# JfxNgs: A BAM/VCF viewer with javascript-based filtering/reformatting functionalities

**DOI:** 10.1101/120196

**Authors:** Pierre Lindenbaum, Matilde Karakachoff, Richard Redon

## Abstract

**Motivation:** Visualizing BAM and VCF files is a common task for biologists, but they’re missing a way to filter and to explore the details of each short-read or variation.

**Results:** In that context, we wrote an interactive java-based interface named *JfxNgs* that uses javascript snippets to filter and reformat BAM and VCF files.

**Availability:** https://github.com/lindenb/jvarkit/blob/master/docs/JfxNgs.md

## 1 Description

Powerful tools like the *Integrative Genomics Viewer* (*IGV*) (Thorvalds *et al*., 2013) are very efficient at visualizing the features laying in a genomic region but they’re not really suitable for exploring the details of each item (quality, functional annotations, …). Biologists in our lab currently explore the VCF files using knime4bio (Lindenbaum *et al*., 2011), a plugin we developped for the knime plateform. But by breaking the VCF structure into a table, we lose important informations like the VCF header, and we cannot use the existing programming interfaces like the ‘*java API for high-throughput sequencing data formats’* (htsjdk) (htsjdk, 2016) to analyze the data. Hence, we have written ‘*JfxNgs*’ a java heavy client displaying the items of some VCF and BAM files. Nevertheles we do not see *JfxNgs* as a competitor of *IGV* but rather like a companion that will be used to get the details of each record in the file.

We have taken advantage that the standard edition of ‘java’ distributed by Oracle contains ‘*nashorn*’, an embedded javascript engine. This engine gives the abality to manipulate java objects using simple javascript statements. When those objects are created by htsjdk (htsjdk, 2016), it gives the users a very simple way to filter or reformat a VCF or a BAM file. We had previously implemented this idea in the tools *FilterVcf* and *FilterSamReads* from the *picard* (Picard, 2017) suite and in *jvarkit* (Lindenbaum, 2015), and we have now implemented it in the *JfxNgs* interface (Fig. 1 and 2).

**Figure 1:**
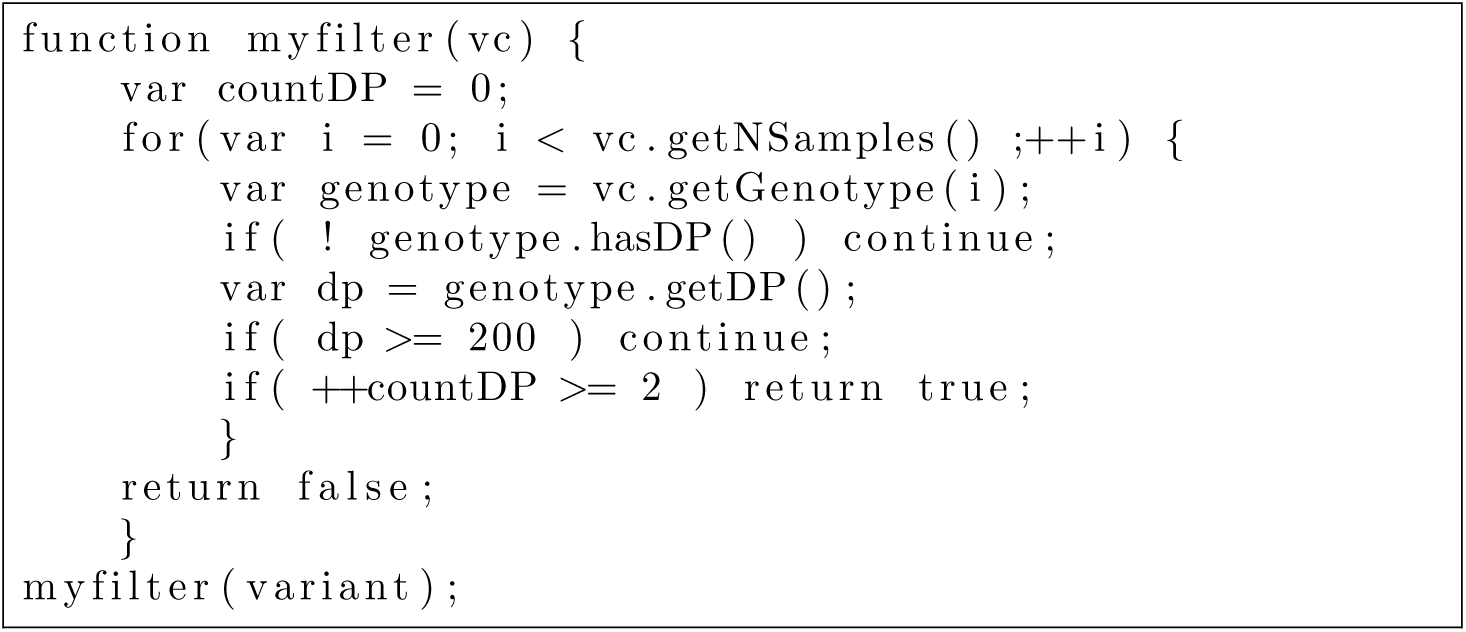
A simple javascript-based filter retaining the variants having at least two genotypes with a depth lower than 200 reads: an instance of the htsjdk java class htsjdk.variant.variantcontext.VariantContext named ‘*variant*’ is injected in the javascript context, the function ‘myfilter’ is invoked. It runs a loop over all the samples and count the number of genotypes having a depth (‘DP’) lower than 200.

**Figure 2:**
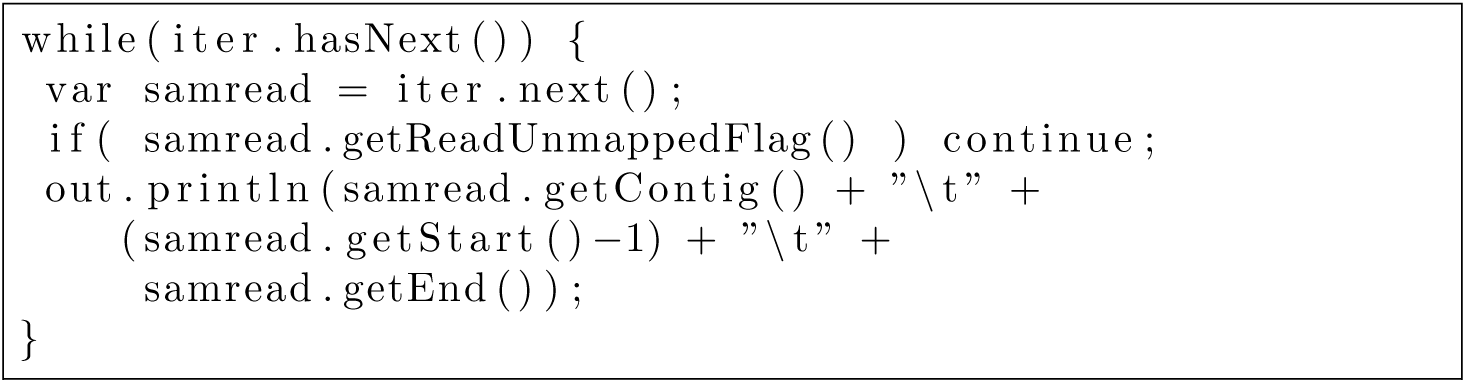
Reformatting a BAM file to a BED file. An iterator named ‘*iter*’, and scanning some instances of the htsjdk java class htsjdk.samtools.SAMRecord, is injected in the javascript context. As long as we can read a SAM record, mapped on the reference, its’ genomic location is printed to ‘*out*’, the current output stream.

**Figure 3:**
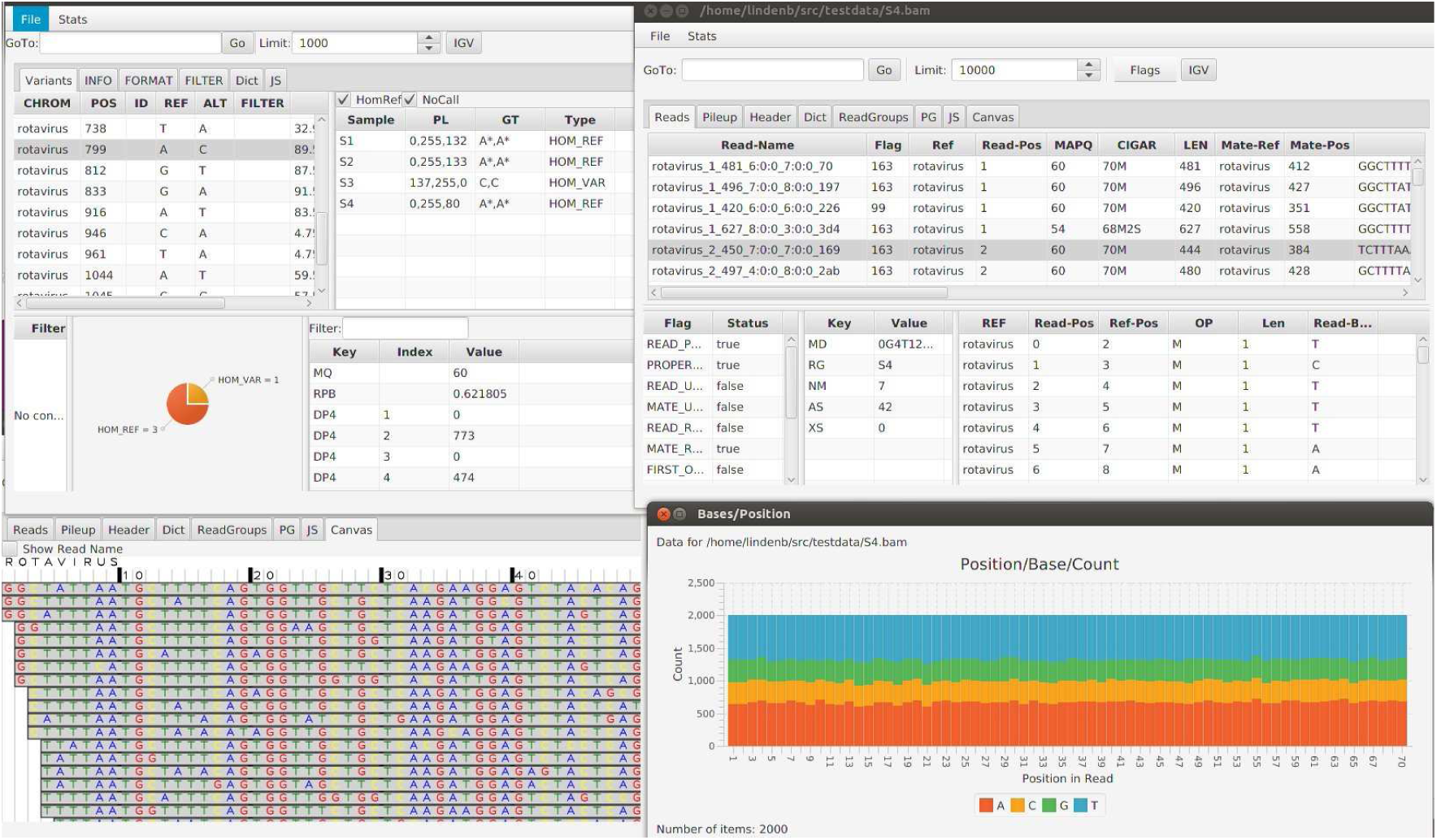
A Screen of jfxngs with a VCF and a BAM window. The ‘JS’ tabs provide a text area to write a javascript code to filter the data.

*JfxNgs* can read local files as long as they’ve been indexed with tabix or tribble. It can also access the remote files if the hosting server supports ‘Byte-Range’ requests. The main window provides tools for indexing BAM and VCF files. The VCF and the BAM windows have common functionalities: filtering the data with javascript, displaying the data in a very simple genome browser, viewing the selected item in a web browser for a common database like Exac (Song *et al*., 2015), displaying simple statistics (Fig. 3), exporting the data with javascript, viewing the current selected genomic interval in *IGV*, viewing the components of the file header, saving the filtered data in a new file. The javascript areas contain a growing library of code snippets to help the user.

The BAM window shows the standard columns of the BAM specification (Read-Name, Sam-Flags, etc…), it provides some tables to view the details of the cigar string, the sam flags, the supplementary alignments. A pileup-like table displays the bases covering each position.

The VCF window displays the standard columns of a VCF file (CHROM, POS, etc…), it provides some tables to clearly view the INFO, FILTER, ALTS data. If the annotations of a prediction algorithm like VEP (McLaren *et al*., 2010) of SNPEff (Cingolani *et al*., 2012) are detected, the predictions for each transcripts are displayed in a new table. A table of each genotypes displays the calls for each variant with the ability to filter out the Hom-Ref and No-Call genotypes. A pedigree file can be associated to a VCF and is then used to detect the Mendelian incompatibilities.

## 2 Availability

The software is available in the jvarkit package (Lindenbaum, 2015). It has been tested under Linux, Windows and MacOS and is freely available at https://github.com/lindenb/jvarkit/blob/master/docs/JfxNgs.md At the time of writing, our tool is also available, packaged as a java webstart application, at http://redonlab.univ-nantes.fr/publimhtml/jnlp/jfxngs, meaning that it doesn’t require any installation but java and an always up-to-date application is downloaded each time the user invokes it.

## Acknowledgements

We want to thank the bioinformatics core facility of Nantes (Biogenouest)for technical support.

